# Spontaneous Intersibling Polymorphism in the Development of Dopaminergic Neuroendocrine Cells in Sea Urchin Larvae: Impacts on the Expansion of Marine Benthic Species

**DOI:** 10.1101/2023.12.03.569790

**Authors:** Alexandra L. Obukhova, Marina Yu. Khabarova, Marina N. Semenova, Viktor V. Starunov, Elena E. Voronezhskaya, Evgeny G. Ivashkin

## Abstract

Plasticity of the nervous system enables the formation of the most adaptive neural circuits and the corresponding behavior of animals. The mechanism by which plasticity arises during development and its involvement in animal adaptation is one of the astonishing questions. Sea urchin larvae are known for their evolutionary and ecological diversity as well as their developmental forms and behavioral patterns. This research addresses the intricate neuroendocrine adaptations that govern larval development of sea urchins, focusing on the coordination between dopaminergic (DA) and serotonergic (5-HT) neurons. The study reveals a heterochronic polymorphism in the appearance of post oral DA neurons and confirms the stable differentiation pattern of apical 5-HT neurons in the larvae of Mesocentrotus nudus and Paracentrotus lividus. We demonstrate that an increased number of DA cells and DA application correlate with downward swimming of the larvae. In contrast, 5-HT cells and serotonin application unsure larval upward swimming. As a result, the 5-HT/DA ratio underlay stage-dependent vertical distribution of the larvae within the water column. In larvae of the same age, the precise balance of 5-HT and DA cells underlie the basis for the different potentials of individuals for upward and downward swimming. This coordination in humoral regulation underlies shifts in larval behavior within a single generation. Based on our findings on DA-cells polymorphism, we have proposed a model illustrating how the balance between the serotonin and dopamine systems, shaped by heterochrony in DA cell appearance, impacts larval behavior, reduces competition between siblings and ensures optimal population expansion. The study explores the evolutionary and ecological implications of these neuroendocrine adaptations in marine species.

## 1 Introduction

The process of development in animals is intricately tied to adaptability, involving mechanisms that maximize reproduction and trade-offs for population stability. Plasticity, or the ability to adjust to different conditions, plays a crucial role at the embryonic development level (DeWitt et al., 1998). Investigating these trends often requires delving into physiological aspects that reveal features not immediately apparent.

One fascinating dimension of adaptability lies in the plasticity of integrated systems within organisms, such as the nervous and endocrine systems (Leroith et al., 1986). These systems collaborate to ensure the overall well-being of the organism, with development representing a series of adjustments animals make to thrive in their environments. These adjustments involve intricate mechanisms, some of which may not be immediately obvious due to their connection to physiology.

The biphasic life cycle, crucial for various aquatic animals (Degnan and Degnan, 2006; Nielsen, 2009, 2012), involves a free-swimming larval phase and a benthic or sessile adult stage. Larval swimming relies on active migration in the planktonic environment, primarily driven by biogenic monoamines such as serotonin (5-HT) and dopamine (DA), orchestrating the coordinated beating of locomotory cilia (Conzelmann et al., 2011). The ciliary beating pattern, determined by the levels of these substances and the number of neurons producing them, varies across species with distinct developmental strategies, underscoring the intricacies of developmental adjustments.

Sea urchins, a classical subject for studying larval swimming patterns, ciliary beating, adaptational polymorphism, and the role of 5-HT and DA in these processes, offer valuable insights (Wada et al., 1997; Adams et al., 2011). During the planktonic phase, larvae exhibit behaviors crucial for survival, including food uptake, distribution, and horizontal migration, all linked to ciliary beating along larval arms. The regulation of ciliary activity is mediated by classical monoamines and other neurotransmitters. Moreover, DA plays an complex role in the sea urchin larval development, influencing locomotion, responses to food, and metamorphosis (Burke, 1983; Wada et al., 1997; Adams et al., 2011). This integrative function of classical neurotransmitters, conserved across various animals, including vertebrates, makes it one of the most enduring neuroendocrine systems in Bilateria’s evolution (D’Aniello et al., 2020).

Both 5-HT and DA neurons play pivotal roles in the regulation of ciliary activity in specialized larval structures known as ciliary bands, which serve as the principal swimming and feeding organs of the larvae. Previous studies have demonstrated that the application of both monoamines stimulates cilia at the early post-hatching stage, while at late gastrula and pluteus stages, the effects tend to become opposite: stimulatory for 5-HT and suppressive for DA (Soliman, 1983; Mogami et al., 1992; Yaguchi and Katow, 2003). Correspondingly, 5-HT increases and DA decreases the beat frequency averaged over the ciliated epithelium, demonstrating stabilized and fluctuate effects on the phases of ciliary stroke (Wada et al., 1997; Shiba et al., 2002). Consequently, the application of various monoamines can initiate retardation or acceleration of swimming speed and modulate the swimming direction in free-swimming sea urchin larvae (Yaguchi and Katow, 2003). Similar effects of monoamines on cilia activity have been demonstrated for the larvae of different animals. External application of 5-HT activates cilia beating and accelerates embryonic rotation within the egg in gastropods (Uhler et al., 2000; Kuang and Goldberg, 2001; Kuang et al., 2002). In gastropod Lymnaea, both 5-HT and DA increase the rate of embryonic rotation with similar potency (Voronezhskaya et al., 1999; Goldberg et al., 2011). In bryozoans Bugula larvae, 5-HT and DA demonstrate opposite actions in the regulation of locomotory-dependent phototaxis (Pires and Woollacott, 1997). Both 5-HT and DA are involved in ciliary activity regulation in vertebrates (Walentek et al., 2014; Zhang et al., 2018).

The sea urchins *Mesocentrotus nudus* and *Paracentrotus lividus* are common species in the Sea of Japan and the Mediterranean Sea, respectively. Both species share similarities in their ecology, being covered with spines and moving slowly along the seabed. These animals rely on swimming planktonic larvae for distribution. After gamete spawning and external fertilization, the embryo hatches at the blastula stage, and the developing larva swims upward to the subsurface layer. During their development, the larvae undergo a transformation, acquiring several pairs of arms and initiating a migration along the water column. This migration is facilitated by both the active beating of cilia in the arms and the influence of natural ocean currents that passively carry the larvae. A competent larva, responding to environmental signals, settles down and undergoes metamorphosis into a new benthic adult. Both *M. nudus* and *P. lividus* have well investigated development, and served as an object in a number of developmental and ecological studies (Agatsuma, 2020; Boudouresque and Verlaque, 2020).

Our research employs a combination of morphological analysis and behavioral assays to explore the coordinated effects of the DA and 5-HT systems on the larval swimming patterns at the gastrula-four arm pluteus stages of two sea urchin species, *Mesocentrotus nudus* and *Paracentrotus lividus*. We have observed that in both species, the stable differentiation pattern of 5-HT-containing neurons is accompanied by significant heterochrony in the appearance of DA-containing cells in larvae at the same developmental stage. These variations in the balance of 5-HT and DA cells underlie the different potentials of same-age larvae for upward and downward swimming along the water column and suggests decreased siblings’ competition. This intricate interplay of neurotransmitters in sea urchin larvae sheds light on their phenotypic plasticity and adaptability. This insight is valuable for understanding how animals adapt to their environments, with potential applications in marine ecology and developmental biology.

## 2 Results

Both *M. nudus* and *P. lividus* have planktotrophic larvae, exhibiting high similarity in larval general morphology throughout their development. Gastrulation is completed approximately 28 hours post-fertilization (hpf). By the prism stage (32 hpf), larvae have developed an intestinal tract comprising the mouth, esophagus, stomach, and anus. The first pair of arms (post-oral arms) begins to form during the prism stage and persists into the early pluteus (36 hpf) and pluteus (40 hpf) stages. Ciliary bands, crucial for larval swimming, develop along the edges of the arms and the oral lobe.

Simultaneously, the apical organ differentiates at the apical pole of the oral lobe during the described stages.

### 2.1 Developmental Features of Post-oral Dopaminergic and Apical Serotonergic Cells in *M. nudus* and *P. lividus*

We utilized the FaGlu method for catecholamine visualization, facilitating the identification of dopamine-containing (DA-containing) cells in intact larvae and cells that uptake dopamine (DA-uptaking) in larvae treated with a low concentration of DA (DA-treated). This method enables detailed visualization of neuronal morphology, including processes. Immunoreactivity against serotonin was employed to visualize 5-HT-containing cells in the larval apical organ.

In intact *M. nudus* larvae, no DA-producing cells were visible in the gastrula and prism stages (28 and 32 hpf) (Fig. 1 A-B). Four DA-producing cells emerged at the base of each post-oral arm at the 36 hpf early pluteus stage (Fig. 1 C) and retains at the 40 hpf pluteus stage (Fig. 1 D). Serotonin-containing cells appeared in the apical organ area, starting with one cell at the gastrula stage (Fig. 1 E) and gradually increasing to 4, 6, and 8 cells during the transition from prism to pluteus (Fig. 1 F-H). Thus, both the number of DA-containing cells in the post-oral region (Fig. 1 M) and 5-HT cells in the apical organ area gradually increase during *M. nudus* larval development.

**Figure 1.**
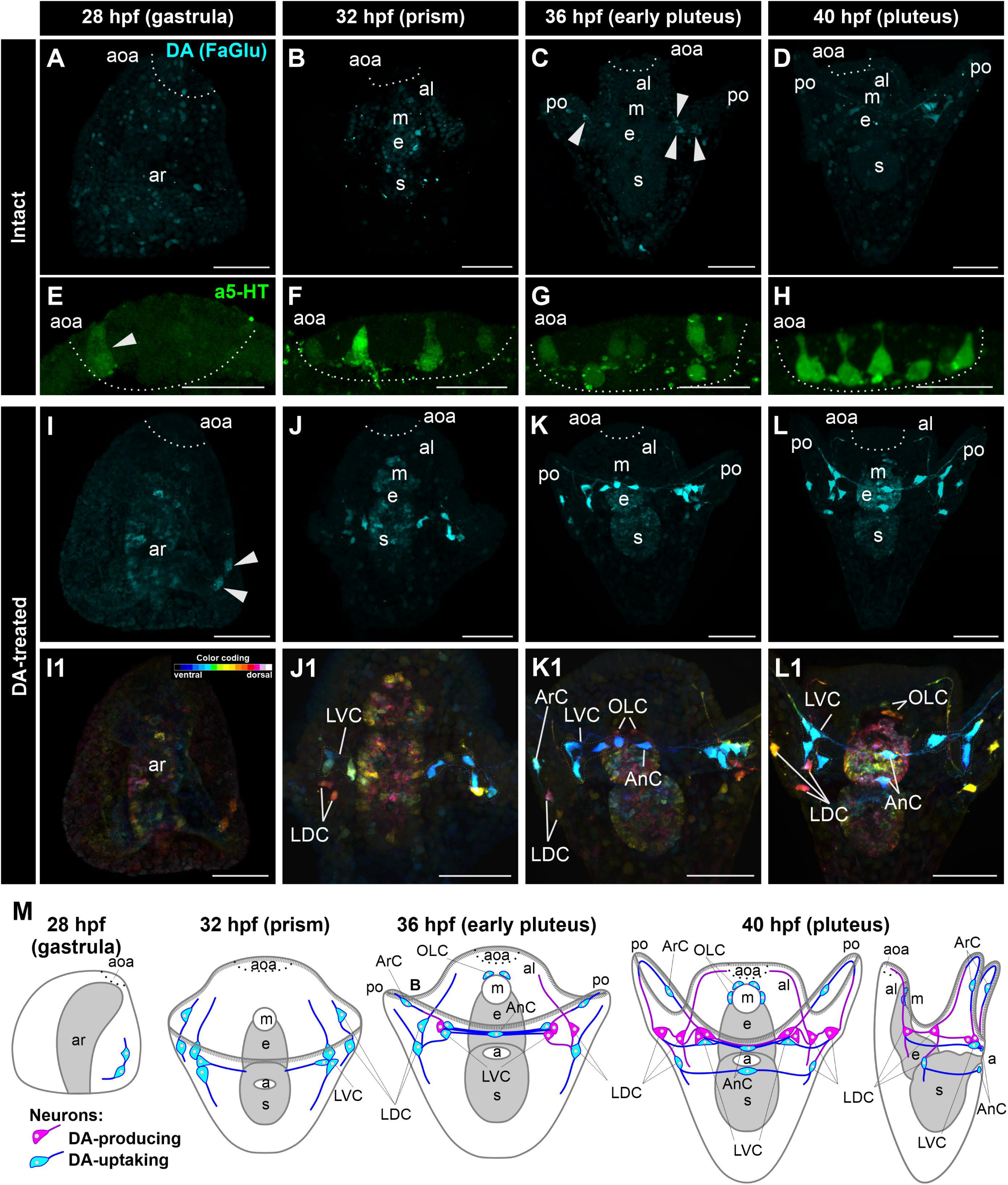
The Developmental Features of DA and 5-HT Neurons in *M. nudus* Larvae. DA-producing cells are visualized in intact larvae, while DA-uptaking cells are shown in DA-treated larvae (FaGlu, cyan). 5-HT cells (a5-HT, green) are located in the apical organ. Panels A-D depict the appearance and gradual increase in DA-producing cells from the gastrula to pluteus stages. The first four cells appear at 36 hpf, and persist at 40 hpf (arrowheads). Panels E-H show the gradual increase in 5-HT cells from one cell at 28 hpf to eight at 40 hpf within the apical organ. Panels I-L display the gradual increase in the number of DA-uptaking cells from two at 28 hpf (arrowheads) to fourteen at 40 hpf. Note association of most post-oral DA cells with the arm edge. DA processes emanate from cell bodies and connect DA cell groups. Panels I1-L1 indicate the relative position of DA-uptaking cells within the larval body, emphasizing the location of LDC group and LVC groups on dorsal and ventral sides, respectively. Panel M provides schematic drawings of DA-producing (magenta) and DA-uptaking (cyan) cells in the development of *M. nudus* larvae from gastrula to pluteus stage. The number of DA-uptaking neurons surpasses that of DA-producing neurons and reaches a maximum at the pluteus stage. Abbreviations: aoa – apical organ area; AnC - unpaired cell near the anus; al – anterior lobe; ar — archenteron; ArC - arm cells; e — esophagus; LDC – lateral dorsal cells; LVC – lateral ventral cells; m — mouth; OLC – oral lobe cells; po — post-oral arms; s — stomach.

In DA-treated *M. nudus* larvae, two DA-uptaking cells were detected on the ventral side at the base of the archenteron as early as gastrula (28 hpf) (Fig.1 I, I1). By the prism stage (32 hpf), three cells condensed in the group at the base of the anterolateral arms (Lateral Ventral Cells, LVC). And the bodies of two cells were located at the base of the oral lobe (Lateral Dorsal Cells, LDC group; Fig.1 J, J1). The DA-positive processes extended along the arm edge and toward each contralateral group. In the early pluteus (36 hpf), solitary cells appeared on the post-oral arms (Arm Cells, ArC), and their processes also passed near the arms edge. The left and right groups at the base of the post-oral arm were connected by several processes, and with the unpaired solitary cell located near the anus (Anal Cells, AnC; Fig. 1 K, K1). Additionally, two cells were located near the mouth opening (Oral Lobe Cells, OLC, Fig. 1 K). At the pluteus stage (40 hpf), the LDC group included three cells, and an unpaired cell was added near the anus (AnC). Most of the cells were triangular-shaped multipolar cells sending projections toward each other (Fig. 1 L, L1). LVC and LDC cell groups were connected by a bundle of processes, most of which ran near the arm edge (Fig. 1 M). Four cells were located near the mouth opening (OLC; Fig. 1 L). Throughout all developmental stages, archenteron cells demonstrated slight staining indicating the ability of archenteron and forming stomach cells to uptake DA (Fig.1 I-L, I1-L1). Thus, the ability of cells to uptake DA appears earlier than their ability to synthesize DA in *M. nudus* larvae development, and the number of DA-uptaking cells prevails over that of DA-containing in the same larval stage (Fig. 1 M). The location of cell bodies and their projections along the arm edge demonstrates the association of all post-oral neurons with arm ciliary bands.

In intact larvae of *P. lividus*, the pattern of DA and 5-HT cell appearance was essentially similar to *M. nudus*. No DA-producing cells were visible at the gastrula and early pluteus stages (24, 32, and 36 hpf), although pluteus larvae already developed post-oral arms (Fig S2 A, B, C, D). Two DA-producing cells appeared at the base of post-oral arms only at the pluteus stage (40 hpf). Their processes were visible along the arm edge. Two cells appeared near the mouth (Fig. S2 E). Simultaneously, serotonergic cells appeared in the region of the apical organ, starting with six cells at the 24 hpf (inset Fig S2 A) and ten cells at the 28 hpf gastrula stage (inset Fig S2 B), with a subsequent increase to twelve at 32 hpf, 36 hpf, and 40 hpf (Fig. S2 insets in C, D, and E).

In DA-treated *P. lividus* larvae, a single DA-uptaking cell was visible at the 24 hpf stage in about 20% of the examined specimens (Fig. S2, F, K, K1), and in some larvae, two neurons were visible at the 28 hpf stage (Fig. S2, G, L, L1). By 32 hpf, three cells were located at the base of each post-oral arm (LVC), and their processes could be traced in the arm edge. Note the marked asymmetry in the appearance of the cells – one cell on the left side and two cells on the right side, and vice versa (Figs. S2, H, H1, M, M1, M2). The number of cells increased steadily and reached eight and fourteen at 36 hpf and 40 hpf pluteus, respectively (Figs. S2, I, N, J, O). The cell bodies were concentrated at the base of post oral arms (LDC, LVC groups, Figs S2, N2) and solitary cells located along the arm egde (ArC, Fig. S2, O2). Cells of the post oral arm groups LVC and LDC connected by processes running along the arm (Figs S2, M1, O1). While in *M. nudus* oral lobe cells concentrated preferentially around the mouth (Fig. 1, K1, L1, M), in *P. lividus* only two OLC cells located close to the mouth and another two situated at the edge of anterior lobe (Fig. S2, O2). No DA-cells were found in the anus region in *P. lividus*.

**Figure 2.**
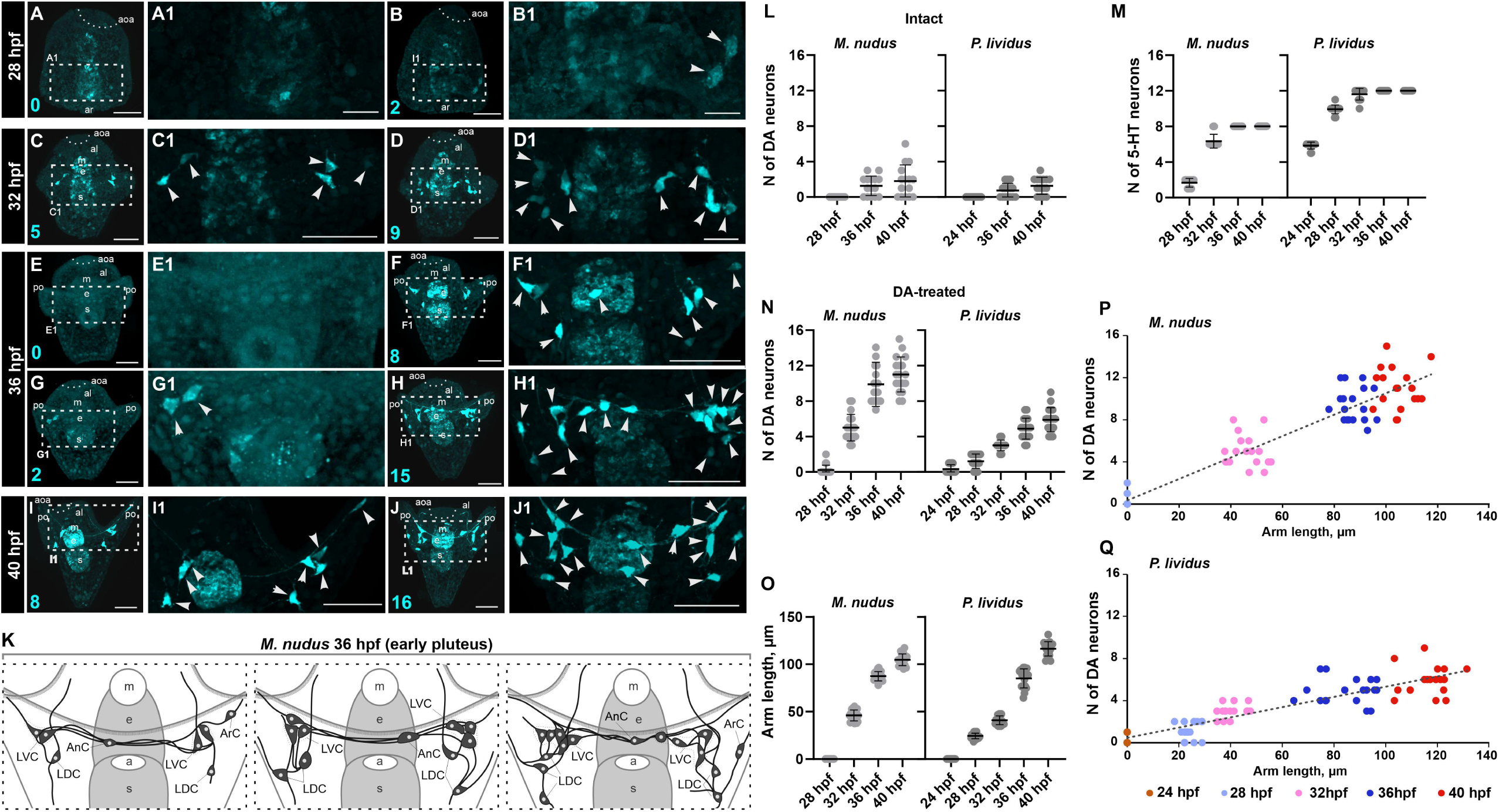
Heterochronic Polymorphism in the Appearance of DA Neurons in Larvae of *M. nudus* and *P. lividus*. Panels (A-J) provide a general view, while panels (A1-J1) offer a high magnification of the neuron-containing region at larvae of subsequent developmental stages from 28 hpf gastrula to 40 hpf pluteus. Both DA-producing and DA-uptaking cells are visualized in cyan. The number of DA neurons is indicated in cyan in panels A-J. Panels A-B1 show variations from zero to two cells in the gastrula stage. Panels C-D1 present samples of variations from five to nine cells at the prism stage. Panels E-H1 depict maximal variations in the number of DA cells from zero to fifteen in the early pluteus stage. Panels I-J show samples of variations from eight to sixteen cells at the pluteus stage. Note processes emanating from DA cells and connecting DA cell groups. Panel K provides schematic drawings of DA cell polymorphism at the early pluteus stage. Note the variation in DA cell number in LDC and LVC groups in one and the same age 36 hpf pluteus larvae. Panels L, M, N, and O present the number of DA-containing neurons, 5-HT-containing neurons, the number of DA-uptaking cells, and arm length at subsequent developmental stages of *M. nudus* and *P. lividus* larvae. Panel P illustrates variations in the number of DA neurons in *M. nudus* according to larval arm length. Panel Q illustrates variations in the number of DA neurons in *P. lividus* according to larval arm length. Mean±SEM are given on graphs. Abbreviations: aoa – apical organ area; al – anterior lobe; ar — archenteron; ArC - arm cells; e — esophagus; m — mouth; po — post-oral arms; s — stomach.

Despite their high similarity, M. nudus and P. lividus larvae exhibit specific features in the timing and number of dopamine (DA) and serotonin (5-HT) neuron appearance. 5-HT neurons differentiate earlier than DA cells in both species. DA-uptaking cells appear earlier in M. nudus than in P. lividus (28 and 32 hpf, respectively), with the total number being lower in P. lividus by 40 hpf. DA-treated larvae in both species exhibit cells capable of uptaking DA, with the number of DA-uptaking cells surpassing that of DA-producing cells at each developmental stage. Notably, cells observed in DA-treated larvae at early stages correspond well with DA-containing cells in intact larvae at later developmental stages. The processes of DA cells in both intact and DA-treated larvae can be traced along the arm edge, indicating that forthcoming DA-producing cells acquire the ability to uptake DA early in differentiation compared to DA synthesis during sea urchin development.

### 2.2 Heterochronic Polymorphism in the Appearance of Dopaminergic Neurons in Larvae

The FaGlu method for dopamine visualization allows efficient and rapid identification of both dopamine-containing (DA-containing) cells in intact larvae and cells that uptake DA (DA-uptaking) in DA-treated larvae, as well as tracing of DA processes. Immunoreactivity against 5-HT is used to visualize 5-HT-containing cells in the larval apical organ. Our methodological approach enables quantitative analysis of DA and 5-HT neurons, allowing correlation with the length of arms at pre-feeding larval stages from gastrula to pluteus. At pre-feeding stages, the arm length precisely reflects the larval age in both *M. nudus* and *P. lividus*.

In *M. nudus* gastrula, the majority of embryos lack any DA cells (Fig. 2A, A1), while some individuals manifest the presence of two DA-uptaking neurons (Fig. 2B, B1). At 32 hpf prism, individuals with five and nine DA-uptaking neurons can be found (Fig. 2 C-D1). The maximum variability in DA-uptaking cells is observed at 36 hpf early pluteus (Fig. 2, E-H1). Some individuals demonstrate the absence or just two DA cells (Fig 2 E, E1, G, G1), while others possess eight and maximally fifteen DA-uptaking cells (Fig 2 F, F1, H, H1). At 40 hpf pluteus stage, all examined individuals have DA cells, with the number varying from eight (Fig. 2 I, I1) to sixteen (Fig. 2 J, J1).

The maximal heterochrony occurs at 36 hpf early pluteus in both DA-uptaking cell number and DA cell variations within different groups of post-oral DA cells. The maximum variations occur in the LDC group located at the base of post-oral arms. This group consists of three to five cells, and some individuals may lack this group completely. The LVC group at the base of the oral lobe varies in number from one to three in different individuals. Anal cells (AnC) appear either as one or two unpaired cells in different individuals. We also noted asymmetry in the appearance and distribution of cells in left and right groups. Heterochronic polymorphism in the presence of DA cells in each group of post-oral cells is summarized in a scheme and expressed as the overall number of DA cells and in the distribution of cells among different post-oral groups (Fig. 2 K). The variation in cell number between individuals is most prominent in pluteus and strongly expressed in LDC and LVC groups.

Analysis of larvae of different ages confirms our observation that within one stage, the number of both DA-containing and DA-uptaking cells varies significantly (Fig. 2 L, M). Within the same age group, larvae exhibit twofold differences (e.g., 28 hpf) and even fourfold differences (36 hpf, 40 hpf) in counts. In the case of DA-containing cells, this variation includes individuals completely lacking DA cells. In the case of DA-uptaking cells, only gastrula larvae lack DA cells, while older larvae possess several DA cells. Such significant differences in the number of DA-uptaking cells in developing larvae are supported by a high variance in their number (Table 1). In contrast, the number of 5-HT cells is stable for a certain larval stage (Fig. 2 M) and demonstrates low variance (Table 1).

**Table 1.**
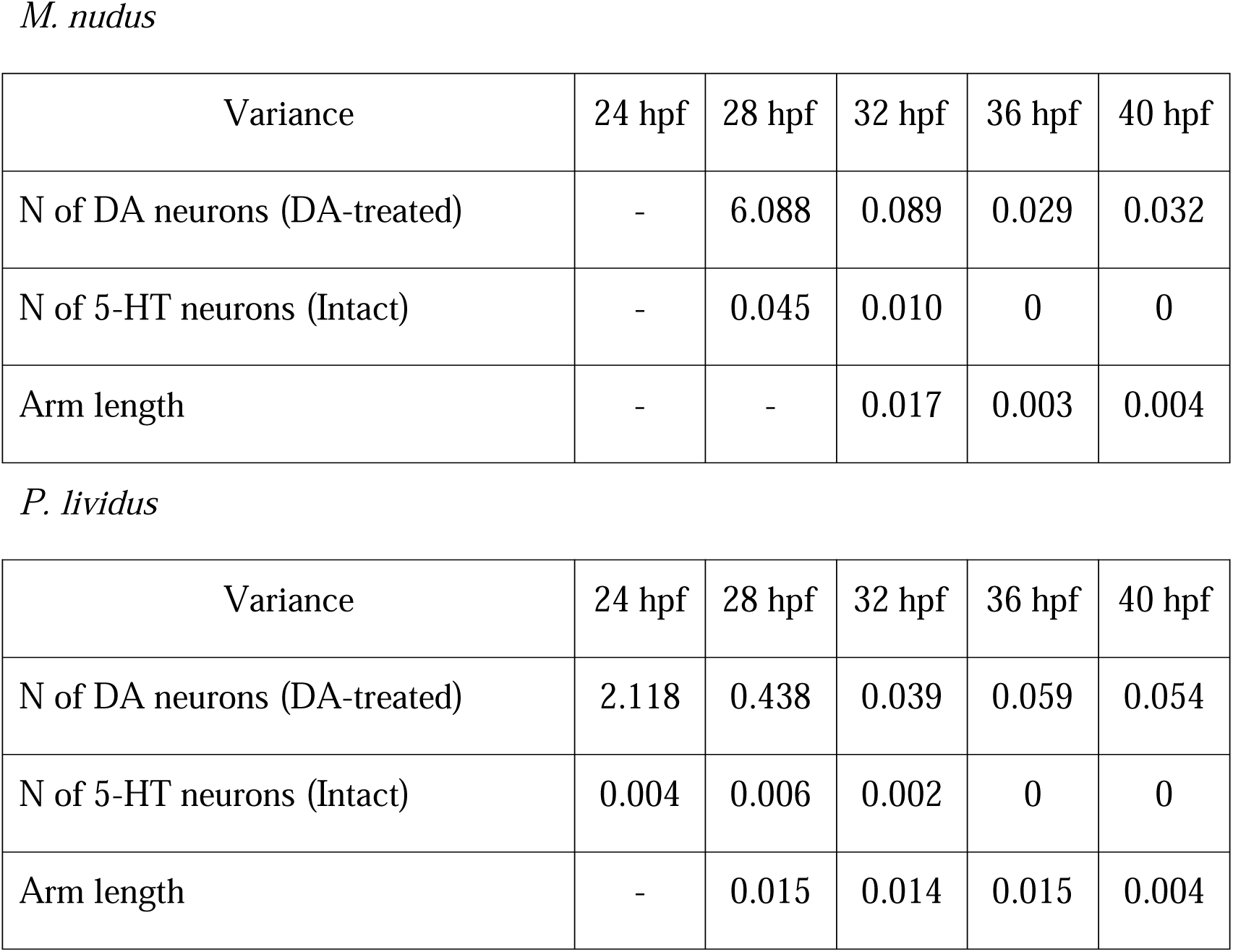
Variances in the Number of 5-HT and DA Neurons in Larvae of *M. nudus* and *P. lividus* at Subsequent Stages of Development.

| Variance | 24 hpf | 28 hpf | 32 hpf | 36 hpf | 40 hpf |
| --- | --- | --- | --- | --- | --- |
| N of DA neurons (DA-treated) | - | 6.088 | 0.089 | 0.029 | 0.032 |
| N of 5-HT neurons (Intact) | - | 0.045 | 0.010 | 0 | 0 |
| Arm length | - | - | 0.017 | 0.003 | 0.004 |

*P. lividus*
| Variance | 24 hpf | 28 hpf | 32 hpf | 36 hpf | 40 hpf |
| --- | --- | --- | --- | --- | --- |
| N of DA neurons (DA-treated) | 2.118 | 0.438 | 0.039 | 0.059 | 0.054 |
| N of 5-HT neurons (Intact) | 0.004 | 0.006 | 0.002 | 0 | 0 |
| Arm length | - | 0.015 | 0.014 | 0.015 | 0.004 |

In *P. lividus* larvae, we observe a similar trend in the appearance of 5-HT and DA cells. While the number of 5-HT cells remains stable for certain larval stages (Fig. 2M, Fig S2, A-E insets, Table 1), DA cells exhibit high variability among individuals of the same age (Fig. S2, I-O2, Table 1). The maximal variations also occurred at the pluteus stage, where individuals with a minimum of three and a maximum of fourteen DA cells can be found (Fig S2, E, O). This observation is supported by neuronal counting at different age larvae (Fig. 2 L, M, M), and high variance in the number of DA neurons (Table 1).

To investigate whether differences in the number of DA cells correlate with larval age, we measured arm length, a characteristic feature of larval developmental tempo. Low variability in arm length was observed among larvae from the same generation (Fig. 2O). Correlation analysis between arm length and the number of DA neurons in selected individuals demonstrates a predicted linear age-related increase in the number of DA cells (Fig 2 P, Q). However, there is no such correlation among larvae of the same age (see Spearman correlation coefficient, Table 2). Individuals with long arms may have a smaller number of DA neurons than larvae with short arms. Such discrepancies are found in all examined developmental stages of *M. nudus* and *P. lividus* (Fig. 2, P, Q, Table 2).

**Table 2.**
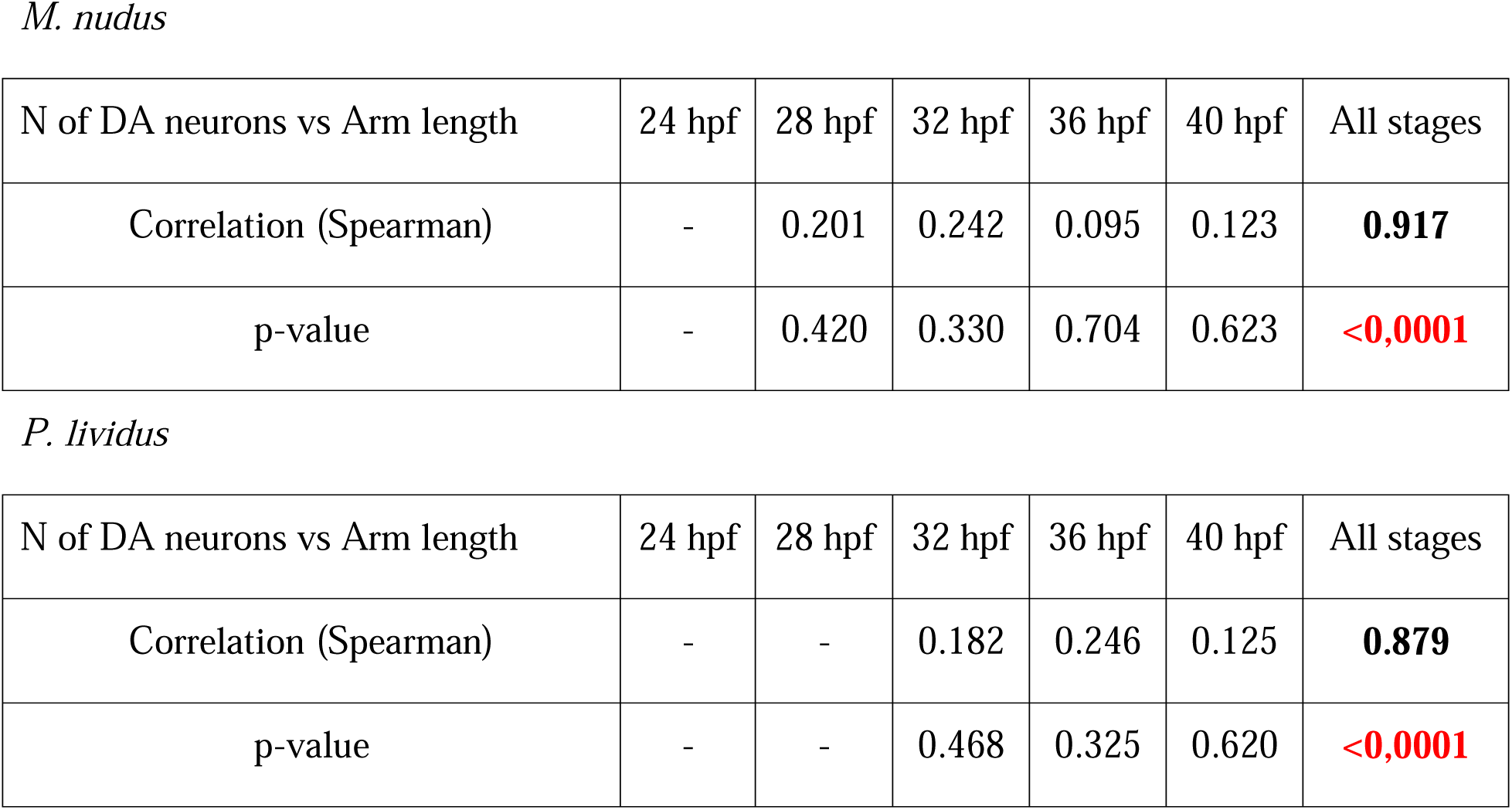
Spearman Correlation Coefficient for Arm Lengths and the Number of DA Neurons in Larvae of *M. nudus* and *P. lividus*.

|  |  |  |  |  |  |  |
| --- | --- | --- | --- | --- | --- | --- |
| N of DA neurons vs Arm length | 24 hpf | 28 hpf | 32 hpf | 36 hpf | 40 hpf | All stages |
| Correlation (Spearman) | - | 0.201 | 0.242 | 0.095 | 0.123 | **0.917** |
| p-value | - | 0.420 | 0.330 | 0.704 | 0.623 | **<0,0001** |
| *P. lividus* |  |  |  |  |  |  |
| N of DA neurons vs Arm length | 24 hpf | 28 hpf | 32 hpf | 36 hpf | 40 hpf | All stages |
| Correlation (Spearman) | - | - | 0.182 | 0.246 | 0.125 | **0.879** |
| p-value | - | - | 0.468 | 0.325 | 0.620 | **<0,0001** |

Thus, our data clearly demonstrate that sea urchin larvae exhibit high polymorphism in DA neuron differentiation and distribution among groups of post-oral cells, almost independently of the general morphology and variations in larval developmental tempo, whereas stage-dependent differentiation of 5-HT apical neurons is highly uniform.

### 2.3 Impact of DA Neuron Number on Larval Downward Swimming Patterns

Our investigation aimed to explore how the heterochrony and the varying number of DA neurons at specific developmental stages influence larval swimming behaviors. Behavioral assays were conducted at three larval stages—24-28 hpf (gastrula), 36 hpf (early pluteus), and 40 hpf (pluteus)— each exhibiting a distinct number of DA neurons. The larvae were tested for swimming ability in 1 m-long test tubes, categorized into three groups: upward-swimming (top), intermediate (middle), and downward-swimming (bottom). We examined the response of larvae at each stage to DA (DA-treated) and 5-HT (5-HT-treated) application (Fig. 3, K). Simultaneously, we counted the number of DA-containing neurons under normal conditions and in the 5-HT-treated groups, along with DA-uptaking cells in the DA-treated group.

**Figure 3.**
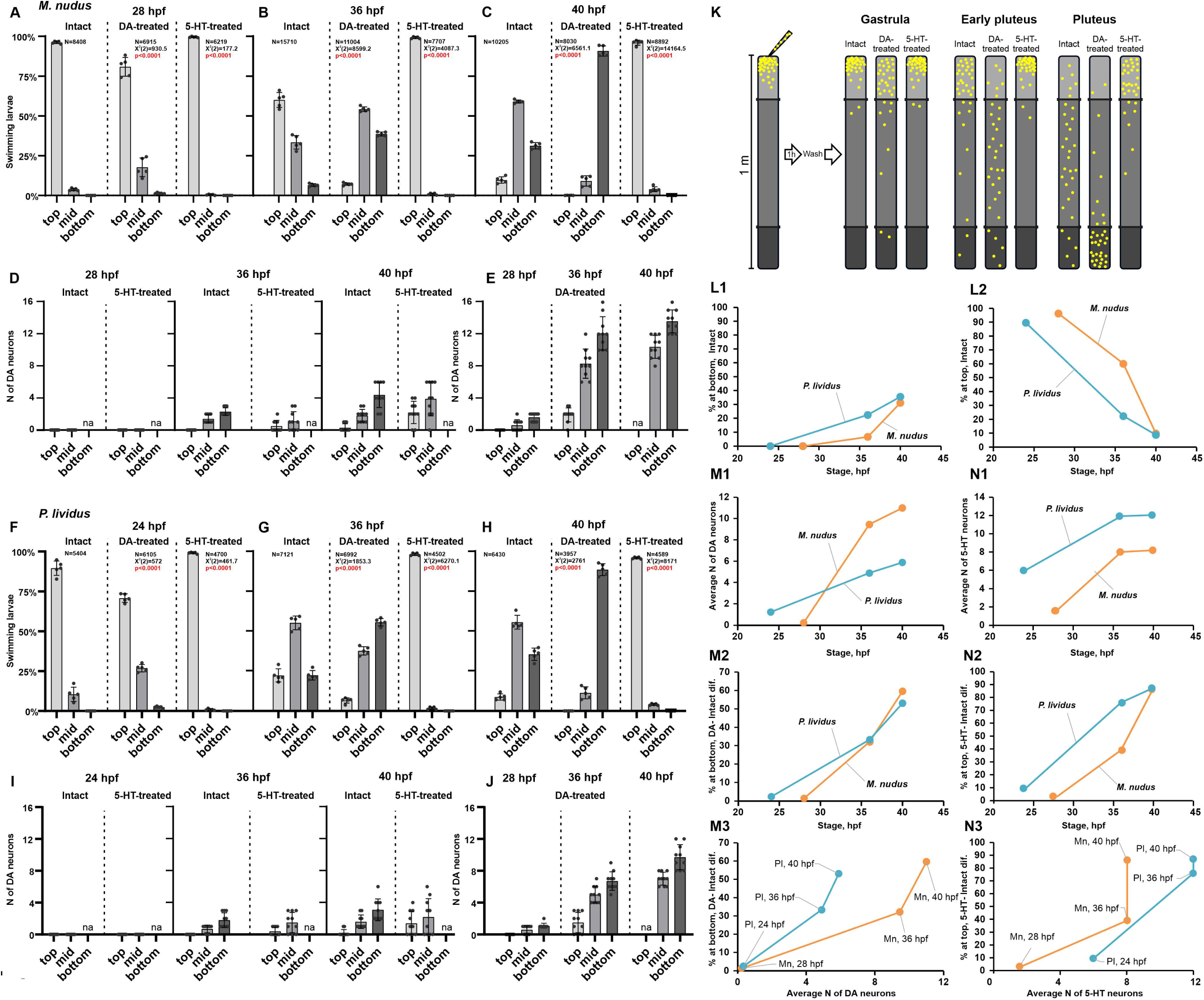
Impact of DA Neuron Count on Larval Swimming Patterns under Normal Conditions and Following DA and 5-HT Application. The relative distribution of larvae at the top, middle, and bottom of the 1 m long water column at subsequent developmental stages is presented for *M. nudus* (panels A-C) and *P. lividus* (panels F-H). Panels D and I show the number of DA-containing neurons, while panels E and J show DA-uptaking neurons related to larvae at various developmental stages and different positions in the water column. Panel K provides a scheme of the experimental setup for the behavioral assay. The number of larvae at the bottom increases (L1), while the number on top decreases (L2) with larval age. Both the number of DA neurons (M1) and 5-HT neurons (N1) increase with larval stages in *M. nudus* and *P. lividus*. There is a positive correlation between the number of larvae at the bottom in the DA-treated group and larval age in both species (M2). Additionally, there is a positive correlation between the number of larvae on top in the 5-HT-treated group and larval age in both species (M2). Furthermore, a positive correlation exists between the number of DA neurons and the relative number of larvae at the bottom in the DA-treated group in *M. nudus* and *P. lividus* larvae of different ages (M3). The correlation between the number of 5-HT neurons and the relative number of larvae on top in the 5-HT-treated group is shown for *M. nudus* and *P. lividus* larvae of different ages (N3). N represents the number of larvae.

In intact 24-28 hpf larvae, the majority exhibited active upward swimming, with only a small percentage in the middle and very few reaching the bottom (intact in Fig. 3, A, F). In both *M. nudus* and *P. lividus*, no DA neurons were found in any larval groups (intact in Fig. 3, D, I). At 36 hpf, about half of *M. nudus* larvae were at the top, one-third in the middle, and approximately 10% at the bottom (intact in Fig. 3, B). In *P. lividus*, half of the 36 hpf larvae were in the middle, and about one-third were at the top and bottom (intact in Fig. 3, G). In both species, the top group lacked DA cells, and a maximum of four DA neurons occurred in the bottom group (intact in Fig. 3 D, I). At 40 hpf, in both species, about half of the larvae were in the middle, one-third at the bottom, and only about 10% remained at the top (intact in Fig. 3 C, H). The maximal number of DA neurons (up to 6 in both species) occurred in the bottom group, with twice fewer cells in the middle group. Larvae from the top group demonstrated the presence of solitary (1-2) DA cells at this stage (intact in Fig. 3 D, I). Thus, in normal conditions, the proportion of the bottom group increased with age, while that of the top group decreased in both species (Fig. 3 L1, L2). This swimming ability correlated with an increased number of DA-containing neurons in older larvae (Fig. 3 M1).

Preincubation with dopamine significantly altered the proportion of downward swimming larvae. At 24-28 hpf, about 25% of the larvae were in the middle and at the bottom (DA-treated in Fig. 3 A, F). Moreover, such individuals demonstrated the presence of DA-uptaking neurons (1-2) even at the gastrula stage (DA-treated in Fig. 3 E, J). At 36 hpf, about half of the *M. nudus* larvae were in the middle, and one-third at the bottom, resembling the swimming pattern of intact but older (40 hpf) larvae of this species (DA-treated in Fig. 3 B, intact Fig. 3 C). Half of the 36 hpf *P. lividus* larvae were at the bottom, and one-third in the middle (DA-treated in Fig. 3 G). In both species, only 10% of the larvae remained at the top. Notably, at this stage, larvae demonstrated maximal variance in the number of DA-uptaking neurons. The minimal number of such DA cells (up to four) occurred in the top group in both species. In *M. nudus*, six to twelve DA cells occurred in the middle group, and a maximum of up to sixteen cells in the bottom group. In *P. lividus*, four to eight cells occurred in medial and bottom groups, with a maximum of ten cells in the bottom group (DA-treated in Fig. 3 E, J). At 40 hpf, most DA-treated larvae appeared at the bottom, with only about 10% in the middle, and a few solitary individuals from the top in both species (DA-treated in Fig. 3 C, H). None or only solitary DA-uptaking cells were detected in 40 hpf larvae from the top group, while DA-containing cells are usually present at the pluteus stage in the intact group. Conversely, the same-stage larvae in the middle group contained eight to ten cells in *M. nudus* and six to eight in *P. lividus*. The maximal number of DA-uptaking cells was found in bottom group larvae of this age and reached sixteen in *M. nudus* and twelve in *P. lividus* (DA-treated in Fig. 3 E, J). Thus, preincubation with dopamine resulted in a significant enhancement in larval downward swimming in both species. Notably, the ability of the larvae to respond to DA became stronger with age.

Preincubation with 5-HT demonstrated the alternative effect to DA. At the gastrula stage, all larvae remained at the top and demonstrated no presence of DA-containing cells (5-HT-treated in Fig. 3, A, F, D, I). Notably, the same swimming pattern, with most of the larvae at the top, occurred for 36 hpf and 40 hpf larvae of both species (5-HT-treated in Fig. 3, B, C, G, H), while the respective individuals contained DA cells similar to intact larvae at the same age (up to four at 36 phf and up to six at 40 hpf). Rare individuals found at the bottom contained no DA cells (5-HT-treated in Fig. 3, D, I). Thus, preincubation with 5-HT caused larvae of all ages to swim upward in both species, and this ability was independent of the number of DA-containing neurons present in the larvae at a particular stage.

Comparing the number of differentiated DA and 5-HT neurons in the larvae of *M. nudus* and *P. lividus* and their tendency to downward and upward swimming at different stages, *P. lividus* larvae generally possessed fewer DA neurons and more 5-HT neurons than *M. nudus* larvae of the same age (Fig. 3 M1, L1). However, in our experiments, both species exhibited approximately equal numbers of downward swimming individuals at each developmental stage (Fig. 3 M2). Even more *M. nudus* 36 hpf larvae appeared at the top than *P. lividus* larvae at the same age. Indeed, our findings reveal that there is no direct correlation between the absolute number of DA neurons and the downward swimming behavior of larvae in interspecies relation. While preincubation with DA consistently influenced larval swimming patterns across different developmental stages, the relationship between the number of DA neurons and swimming behavior appears to be more complex and multifaceted.

Individuals at the bottom may have from 4 (36 hph) to 6 (40 hpf) DA neurons as in *P. lividus*, and from 10 (36 hpf) to 12 (40 hpf) as in *M. nudus* (Fig. 3 M2). The number of 5-HT neurons exhibits a positive correlation with the larva’s ability to swim upward. At 36 hpf, significantly fewer *M. nudus* larvae with a mean of 8 5-HT cells appear at the top than 36 hpf *P. lividus* larvae with a mean of 12 5-HT cells. Notably, by 40 hpf, the majority of larvae from both species predominantly occupy the top portion. (Fig. 3 N3). The number of DA neurons in downward swimming larvae probably does not reflect any ecological specificity but rather is a species-specific characteristic feature. Overall, in all cases, the larvae possessing more DA neurons demonstrated a stronger response to DA application and tended to downward swimming. Thus, we can declare that the number of DA neurons can be treated as equivalent to the position of the larvae in the water column.

## 3 Discussion

This study investigates the presence of post-oral neurons containing dopamine (DA) in sea urchin larvae, specifically *M. nudus* and *P. lividus*. Our findings emphasize the detectability of these neurons throughout development until the initiation of DA expression, highlighting their ability to uptake and store exogenous DA. The incorporation of the FaGlu express method for DA detection facilitated a crucial interspecies comparison. During normal sea urchin larval development, significant heterochrony was observed in both the quantity and timing of these neurons’ appearance in both studied sea urchin species. Furthermore, we identified a correlation between the vertical distribution of larvae in the water column and the number of DA and 5-HT neurons. Our exploration reveals similarities in both heterochrony and monoamine effects in both species, while also noting some species-specific features in terms of neuron quantity, vertical distribution, and the larva’s responsiveness to DA and 5-HT.

### 3.1 Comparison of Dopaminergic Elements in Sea Urchin Larvae

The presence of dopamine (DA) cells has been demonstrated at the four-arm pluteus stage in sea urchins *Psammechinus miliaris*, *Strongylocentrotus droebachiensis*, and *Echinocardium cordatum* (Ryberg, 1974; Bisgrove and Burke, 1987; Nezlin and Yushin, 1994). Authors utilized either histochemical techniques for catecholamine visualization or immunochemical detection of dopamine. Immunoreactivity against dopamine and histochemical fluorescence detection of catecholamine-containing cells revealed identical DA cell bodies and processes in *S. droebachiensis* (Bisgrove and Burke, 1987). Post-oral neurons also exhibited immunoreactivity to the essential enzyme for DA synthesis, tyrosine hydroxylase (TH), in *Lytechinus variegatus* (Slota and McClay, 2018), and aromatic L-amino acid decarboxylase and TH in *Strongylocentrotus purpuratus* (Paganos et al., 2021). Regardless of the visualization methods used in all mentioned species, DA cells were found concentrated in post-oral ganglia or groups of post-oral neurons, later appearing along the base of the arm and circumoral ciliary bands.

In the described sea urchin species *M. nudus* and *P. lividus*, we also detected dopaminergic cells associated with arms. Using histochemical techniques for catecholamine visualization, we identified two groups of cells concentrated at the base of post-oral arms (latero-dorsal group, LDC) and oral lobe (latero-ventral group, LVC). Moreover, we demonstrated that these cells are capable of uptaking dopamine as early as the gastrula stage. This suggests that these dopaminergic cells have already reached a differentiated state and express the DA transporter (an important part of the DA phenotype) earlier than they are able to synthesize dopamine. LDC and LVC groups of DA-uptaking neurons found in *M. nudus* and *P. lividus* correspond well with the post-oral neurons and lateral ganglion neurons described in other species of sea urchin larvae (Bisgrove and Burke, 1987; Slota and McClay, 2018; Paganos et al., 2021). Solitary cells situated in the anus and mouth area emerge subsequent to arm-associated dopaminergic cells in *M. nudus* and *P. lividus*, corresponding to the peripheral DA-containing cells in those regions (Bisgrove and Burke, 1987). Our results allow us to distinguish early differentiating post-oral groups of DA cells and late differentiating cells in the mouth and anus regions in the developing larvae of *M. nudus* and *P. lividus*.

Dopaminergic cells associated with post-oral arms were found in larvae of different sea urchin species (Chen and Adams, 2022). In *Eucidaris tribuloides*, *Arbacia punctulata*, *Lytechinus pictus*, and *Lytechinus variegatus*, dopaminergic groups of cells present at pre-feeding gastrula and pluteus stages resemble LDC and LVC in *M. nudus* and *P. lividus*. At later feeding stages of these species, additional dopaminergic cells appear in the mouth and stomach region. Post-oral DA cells located at the base of post-oral arms, described in the pluteus of *L. variegatus* (Slota and McClay, 2018), and *S. purpuratus* (Paganos et al., 2021), align with the position of the cell bodies and cell projections along the arm edge, corresponding to the morphology of DA-containing and DA-uptaking cells in the corresponding areas of *M. nudus* and *P. lividus* pluteus. However, only late differentiating groups of DA cells in the mouth and stomach regions were present in *Echinarachnius parma*, *Dendraster excentricus*, *Encope michelini*, and *Echinometra lucunter* sea urchin larvae. The authors highlighted the interspecies variation in the number of DA cells, attributing this difference to the phenotypic plasticity of the post-oral arms (Chen and Adams, 2022). Nevertheless, they did not provide data concerning the number of DA cells at comparable developmental stages for each species, and they also did not address potential individual variations within the described species.

In our study, we demonstrated significant variations among individuals of the same species, which remained consistent for both *M. nudus* and *P. lividus* throughout all described stages from gastrula to 40 hpf pluteus. Notably, individuals lacking DA-containing cells coexisted with those containing 2 or even 6 DA cells. Additionally, the absence of cell appearance around the mouth prior to the differentiation of post-oral group cells was consistently observed. The observed differences in the distribution and timing of DA cell appearance within larvae of the same species prevailed over variations in DA cell counts across different species (Chen and Adams, 2022). Our findings underscore the considerable variability in the morphology and timing of DA neurons among individuals within specific sea urchin species. The specific cell number within a particular group may not serve as a stable characteristic feature for the species. Importantly, in our experiments, all larvae were maintained under equal conditions and were not exposed to external stimuli. The identified variation in cell number suggests the presence of environmental-signal independent polymorphism in sea urchin larvae, arising from natural heterochrony in the development of the dopaminergic system. The expression of individual polymorphism in *M. nudus* and *P. lividus* equals or exceeds the interspecies differences described earlier (Chen and Adams, 2022). This underscores the need to consider such individual variations for the accurate interpretation of experimental work and evolutionary-developmental insights.

### 3.2 Post-oral Dopaminergic Neurons Heterochrony and Dopamine-Mediated Larval Polymorphism in Sea Urchins

The preeminent form of polymorphism observed in sea urchins pertains to the varied post-oral arm lengths exhibited by pluteus larvae (Boidron-Metairon, 1988; Strathmann et al., 1992). This phenomenon ensues in response to nutritional cues and chemical determinants of predators (Miner, 2007; Barnes and Allen, 2023). The signaling transpires during pre-feeding larval stages and manifests in subsequent development. Previous investigations have demonstrated that DA signaling, mediated through a type-D2 receptor, governs this type of developmental plasticity (Adams et al., 2011). A correlation is observed, wherein heightened DA levels correspond to diminished arm length. Concurrently, it has been observed that the responsiveness to DA undergoes modifications with larval maturation (Kalachev and Tankovich, 2023). Noteworthy is the existence of analogous data for *M. nudus*, a species explored in this study (Kalachev, 2020).

Our study reveals that, even under standardized feeding conditions and in the absence of algae or predators during the initial phases of larval development, an inherent polymorphism emerges in the quantity of dopaminergic neurons, which serve as the primary source of DA signaling in ontogeny. Consequently, despite the lack of external environmental influences, the examined sea urchin larvae exhibit diversity in DA signaling levels.

Across each investigated developmental stage in both species, there is no statistically significant correlation between the quantity of dopaminergic neurons and the length of post-oral arms. This lack of correlation suggests that the observed polymorphism in neuron count does not manifest as a modulation of arm growth during the pre-feeding larval stages. It implies that the regulatory framework governing arm growth is likely more intricately structured than a direct influence of DA. Alternatively, the rate of DA release may be independently regulated, regardless of cellular quantity. This finding emphasizes the need for a more nuanced examination in future research to unravel the complexities of the regulatory mechanisms at play.

### 3.3 Heterochrony in the Appearance of Post-Oral Dopaminergic Neurons and Larval Swimming Patterns

Earlier, the interscpecies heterochrony in appearance of 5-HT neurons has been described between the larvae of planctotrophic and lecitotrophic sea urchin species (Bisgrove and Raff, 1989). The alteration in the time of nervous system development between direct and indirect developers, implicates heterochronies in cellular differentiation as an important component of the adaptation. Generally, heterochrony, as an evolutionary shift in the relative time of certain structure formations, is widely distributed among developmental programs and serves as one of the sources for evolutionary changes (Kaufman and Raff, 1983; Gould, 1985; Smith, 2003), especially in echinoderms (Parks et al., 1988; Bisgrove and Raff, 1989). Our finding of adaptational heterochrony in nervous system development within the sibling larvae exemplifies neuronal plasticity, potentially underlying evolutionary adaptations within a particular species. The possible cellular mechanism of this phenomenon requires further investigation.

We observed a correlation between the number of DA and 5-HT cells in larvae and their swimming pattern, driven by ciliary beating. Both DA and 5-HT are well-established regulators of cilia activity, and their roles in the control of sea urchin larval swimming have been extensively studied. Previous research has demonstrated opposing effects of 5-HT and DA on the ciliary beating rate of arm bands’ cilia, with 5-HT reducing the beat period and DA prolonging it (Wada et al., 1997). Our experimental results, revealing a positive correlation between the differentiation of DA elements and larval downward swimming ability, are in line with existing data. A higher number of DA-containing neurons potentially can synthesize and release more DA, leading to a reduction in ciliary beating rate. Consequently, larvae lose their swimming ability and sink. 5-HT counteracts this downward swimming by increasing swimming ability through elevated ciliary beating (5-HT reduces the beat period of cilia). Therefore, the balance between the DA and 5-HT systems during development can determine the prevailing position of the larva in the water column.

Our findings indicate that heterochrony in the appearance of DA cells in sea urchin larvae of the same age significantly influences their vertical distribution within the water column. The development of more DA-synthesizing neurons enhances the larva’s ability to sink to the bottom. In contrast, 5-HT is associated with upward swimming, and the quantity of 5-HT cells determines the upper position of the larva. Consequently, in one generation, a limited number of larvae with preferential differentiation of the DA neurons and respective downward ability appear at each developmental stage. We propose that the described synchrony of 5-HT neurons differentiation, together with heterochrony in DA neurons’ appearance, serves as a basis for the optimal vertical distribution of sibling animals, thereby reducing the impact of sibling competition. The possible model of optimal expansion of sea urchin species based on gradual increases in larvae with downward ability is presented in Figure 4.

**Figure 4.**
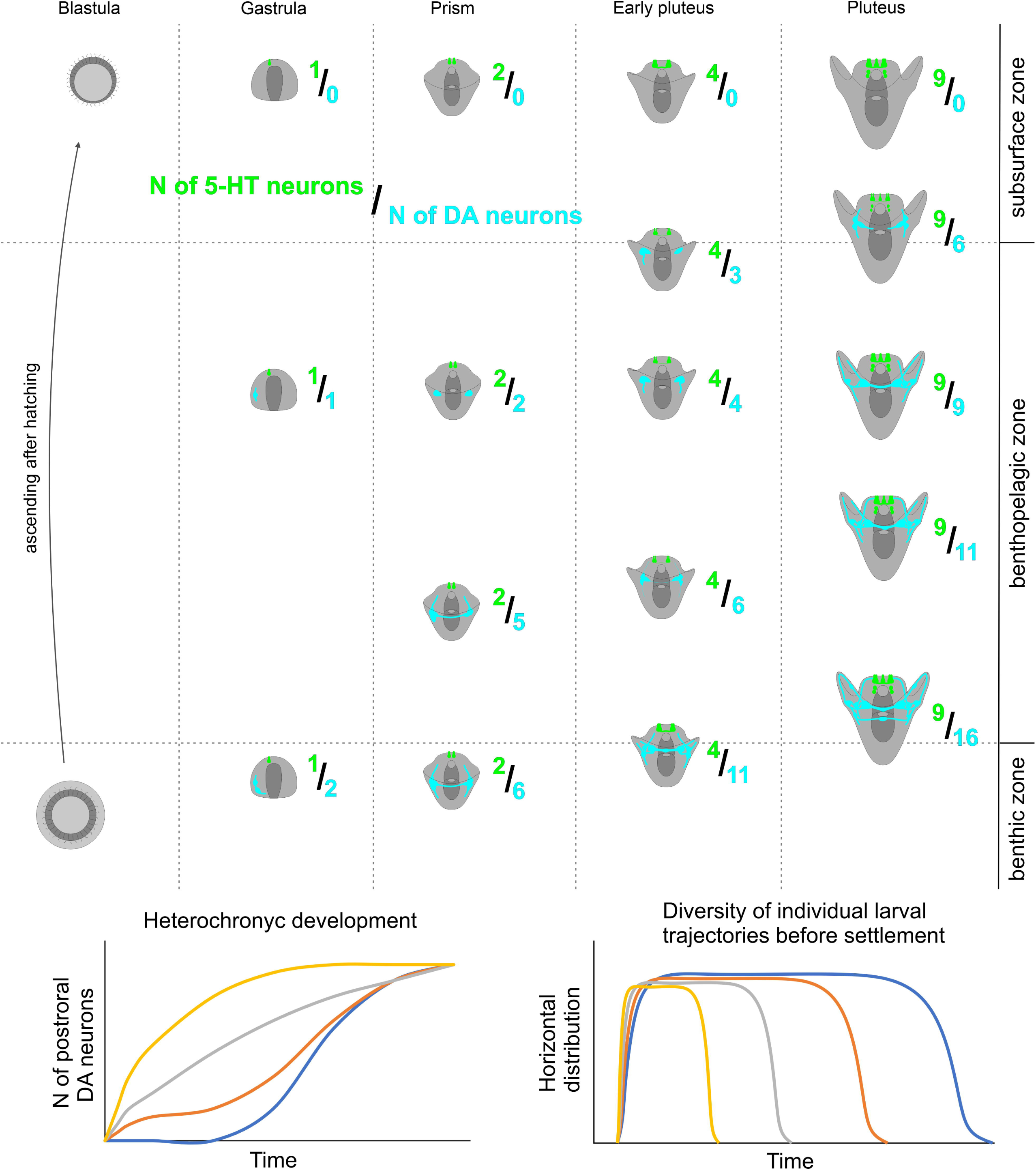
A Proposed Model for Heterochronical Development of Larval Dopaminergic Neurons and Its Impact on Individual Swimming Direction in Sibling Echinoderm Larvae. Generally, the number of both DA and 5-HT neurons increases with the larval age. However, the 5HT/DA ratio exhibits considerable variation in larvae of the same age due to the high heterochronic polymorphism in the number of DA cells. Larvae with a higher count of DA neurons tend to be situated deeper in the water column. This coordination of serotonin and dopamine humoral regulation underlies a shift in larval behavior within a single generation, reducing siblings’ concurrence, and ensuring better larval dispersal and population expansion.

The ability of planktonic larvae to disperse is a crucial factor for the ecological niche of a species (Alexandridis et al., 2017). Planktonic larvae swimming in the upper layers of water can be carried by sea currents for long distances, and the longer the larva remains an active swimmer, the further it can be carried away from the residence of the benthic parental specimens (Jablonski and Lutz, 1983). In temperate sea urchin species, this planktonic stage generally lasts from 2 to 6 weeks (Thorson, 1961), and during this period, an ocean current of only 0.5 km/h could carry a larva for 150-550 km (Scheltema, 1977). Larvae of some tropical species can remain in the plankton up to 6 months or more, traversing trans-oceanic distances (Edmunds, 1977; Scheltema, 1977). On the other hand, some polar and deep-sea species demonstrate demersal larvae that swim or crawl in the near-bottom water layer until settlement (Pearse, 1969). In high latitudes, this dispersal mode appears to be an adaptive compromise, retaining some of the advantages of free-swimming larvae while reducing mortality by keeping larvae out of dangerous surface waters (Pearse, 1969). Also, benthic dispersal at any latitude would expose larvae to a more stable benthic food resource and a temperature-salinity regime compared to the seasonally and diurnally more variable conditions at the sea surface, particularly in nearshore waters (Levinton, 1974; Hendler, 1977; McCall, 1978). The described ability of larvae from one generation and the same age to gradually swim downward and remain near the bottom can be an adaptive mechanism for a gradual change from planktonic development (more efficient for species expansion but more dangerous) to demersal development (not as effective for species distribution but increasing larval survival). As we demonstrated, the change from planktonic to demersal developmental mode occurs within one generation, representing the most effective species strategy to compromise between expansion, feeding, and survival.

### 3.4 Post-oral Dopaminergic Neurons in Sea Urchin, and the Evolution of Endocrine System

Recent studies have revealed that post-oral dopaminergic neurons express the enzyme for acetylcholine synthesis and a cocktail of neuropeptides (Slota and McClay, 2018; Paganos et al., 2021). This demonstrates that post-oral neurons can influence ciliary beating in a more intricate manner than merely through the frequency of beats, determining finer patterns of larval behavior. It happens via neurotransmitter receptors that are located directly located on the cilia (Katow et al., 2010). Moreover, processes from serotoninergic and dopaminergic neurons do not directly extend to each ciliary cell, forming synapses. Thus, 5-HT and DA act on cilia rather as humoral endocrine regulators.

Hypotheses about the origin of pancreatic β cells from neurons have been proposed earlier based on the proximity of their neurogenic marker expression profile (Eberhard, 2013). Recent studies have shown that DA-positive post-oral neurons in sea urchins express Pdx1 (one of the master-regulators for β cells differentiation) and a set of markers that indicate their homology with β cells in the mammalian pancreas (Perillo et al., 2018; Paganos et al., 2021). These findings have sparked discussions about the evolution of the endocrine system and its conservation across different animal phyla.

Our findings suggest the presence of a regulatory mechanism governing the number of post-oral neurons during the development of sea urchins. Typically, the regulation through cell number and mass is more characteristic of endocrine organs, while adaptive physiological regulations through intricate cell interactions are more typical for the nervous system (Leroith et al., 1986). Pancreatic β cells adhere to this pattern, featuring a complex system for controlling cell mass (Bouwens and Rooman, 2005). Consequently, this endocrine organ is distinguished by the determination of developmental trajectories, influencing various aspects of feeding behavior and adaptive physiological traits in adult life (Kim et al., 2010; Castell et al., 2022). Our discovery reveals that in sea urchins, the post-oral neurons is a part of a complex neuroendocrine regulator, resembling its operation in vertebrates. Notably, the natural polymorphism in the number of post-oral neurons determine feeding behavior-related aspects of larval development. This implies that this system was present at the level of the common ancestor of ambulacrarians and chordates in its fully functional form, rather than as distinct components. Moreover, in addition to cellular type homology, there is a direct functional continuity observed in this system between sea urchins and vertebrates.

Our study primarily focuses on serotoninergic and dopaminergic cells as humoral regulators. In vertebrates, 5-HT serves a partially direct endocrine function, while DA has largely relinquished this role over the course of evolution. However, the regulation of cell mass in pancreatic β cells is still intertwined with these ancient monoaminergic regulatory systems. β cells express 5-HT receptors, and 5-HT increases their proliferation in pubertal islets (Castell et al., 2022). DA synthesis is absent in β cells in vertebrates. Nevertheless, DA is synthesized by α cells, while β cells express the DA D2 receptor, acting as a positive regulator of the cell mass of β cells (Sakano et al., 2016). Thus, despite their specialization in neurotransmitter function, monoamines within this endocrine system retain their ancient, overarching integrative role, governing the intricate interplay of feeding behavior and physiological processes at the population adaptation level.

### 3.5 Conclusion and Future Directions

In conclusion our research not only advances the understanding of sea urchin larval development but also contributes to broader discussions on the evolutionary and ecological implications of neuroendocrine adaptations. The intricate coordination between the 5-HT and DA systems, along with the observed shifts in larval behavior within a single generation, underscores the adaptability of these organisms to optimize ecological success. Future investigations should delve deeper into the cellular and molecular mechanisms orchestrating the phenomenon of spontaneous intersibling polymorphism in DA cells, providing a more nuanced understanding of the evolutionary strategies at play.

## 4 Materials and Methods

### 4.1 Animal Handling and Larval Culture

Adult *Mesocentrotus nudus* (A. Agassiz, 1864) were collected in Peter the Great Bay in the Sea of Japan (“Vostok” Biological Station, Zhirmunsky Institute of Marine Biology of Far East Branch of RAS). Adult *Paracentrotus lividus* (Lamarck, 1816) were collected in the Mediterranean Sea near Paphos, Cyprus, and in the marine station of Endoume, France (Station Marine d’Endoume IMBE, Marseille, France). Spawning was induced by intracoelomic injection of 0.5 M KCl. Eggs were collected and fertilized in filtered seawater (FSW). Embryos and larvae were cultured at 18 °C until they reached the gastrula, prism, two-armed early pluteus, and pluteus stages and were used for morphological studies and swimming experiments. Spawned adults and larvae not used for experiments were returned to their natural environment.

### 4.2 Histochemical Detection of DA-Positive Cells

For the histochemical detection of catecholamines (DA, particularly), we used a mixture of 0.5% glutaraldehyde and 4% paraformaldehyde in FSW with 30% sucrose (FaGlu). Samples stored in FaGlu mixture at 4 °C during from one day to two one month until examination. Prior imaging, samples were posted on glass slides, excess FaGlu was thoroughly drawn off, and glass slides were stored in a dry, dark place at room temperature for 12 h for drying. After drying, the samples were coated with paraffin oil and covered with a coverslip. Samples were examined using Leica TCS SP5 (Leica, Germany) confocal laser scanning microscope at 405 nm excitation and 475-485 emission filters (Zoological Institute RAS and Saint-Petersburg State University, Saint Petersburg, Russia). Such a combination of excitation and emission filters allows the specific detection of dopamine-derivative fluorescent products of the reaction (Furness et al., 1977).

### 4.3 Immunohistochemical Detection of Serotonin-Containing Cells

To identify cells containing serotonin (5-HT), larvae were fixed in 4% paraformaldehyde in FSW at the appropriate developmental stages. Prior to antibody treatment, embryos were washed with 0.01 M phosphate-buffered saline pH 7.4 with 0.5% Triton X-100 (PBS-TX) three times, and then incubated for 12–24 h at 4 °C with rabbit antibodies against 5-HT (rabbit, Immunostar, Cat #20080) diluted 1:1000 in 0.5% PBS-TX. After incubation with the primary antibodies, the samples were washed with 0.1% PBS-TX three times and treated with Alexa Fluor 488-tagged goat-anti-rabbit IgG antibodies (Thermo Fisher, Cat # A-11008), diluted 1:800 in 0.1% PBS-TX for 6–12 h at 4 °C. Samples were then washed with PBS three times and immersed in 70% glycerol. The samples were examined under a confocal laser scanning microscope Leica TCS SP5 (Leica, Germany) with the appropriate wavelength filter configuration setting (IDB RAS, Moscow, Russia). Images were analyzed with Fiji software (Schindelin et al., 2012).

### 4.4 Swimming Assays and Analysis

Incubation was carried out in plastic Petri dishes with a diameter of 4 cm. 5-HT or DA (0.1 μM – 1 mM final concentration) was added to petri dishes and allowed to incubate for 30 min. FSW was used for the control group. In the preliminary experiments, a setup according to the previously described (Yaguchi and Katow, 2003) was used. The setup was assembled from two plastic tubes (1 cm diameter and 5 cm in length) placed one above with a narrow channel (diameter 1 mm) between upper and lower parts. The lower tube and the channel between tubes were filled with FSW. 2 mL of the incubation solution containing larvae from Petri dishes was transferred to the upper tubes, and positions were set in a dark place for 1 h. After 1 h swimming assay, the upper and lower parts of the setup were separated, and water with larvae transferred in the 9 cm Petri dishes. The number of the larvae in the upper and lower portions was counted under a dissecting microscope. All experimental procedures with larvae were carried out at 18 °C. The experimental setup is present in Fig S1 A. The number of the downward swimming larvae increased with larval age from gastrula till pluteus for both species (Fig S1 B, C, E-H). *M. nudus* early pluteus demonstrates dose-dependent downward swimming reaction in response to DA application (Fig. S1 D). DA and 5-HT at 1 μM concentrations were selected as the minimum required to produce an effect at the base of these preliminary experiments.

For the swimming experiments, we designed and assembled an experimental setup consisting of several vertical columns of 5 cm in diameter and 1 m high. Each column was a polyethylene soft tube with a closed bottom and open upper parts. Before experiments, tubes were filled with FSW and suspended vertically in a dark place. The experimental larvae from respective groups (DA-treated and 5-HT-treated) were washed in FSW and gently transferred into the upper part of the column. After 1 h swimming assay, the soft plastic tubes were clamped in two places: 25 cm from the top, 25 cm from the bottom, thus separating water column into three parts: top, middle and bottom (Fig. 3, K). The water with larvae was subsequently collected from upper, middle, and bottom parts and transferred to the 0.5 L glass. Larvae were prefixed with the addition of a few drops of 4% PFA. Immobilized larvae were collected from the glass bottom, transferred to the 9 cm Petri dishes, and counted under a dissecting microscope. Simultaneous and quick collection of larvae from three parts of the column made it possible to establish the most accurate percentage of the three types of swimming in the larvae of one stage: top, middle, and bottom. After the calculations, a representative larva from each compartment of each column was processed for the detection of catecholamines as described above. Each experiment was repeated 5 times. All swimming essays were conducted in the dark, to exclude the influence of illumination on the larvae (Yaguchi et al., 2022). The described setup allowed the experimental conditions to be as close as possible to natural environmental conditions: due to the relatively wide diameter of columns and high altitude, the effect of capillary force was minimized, and the penetration of larvae into any part of the column could occur only due to the movement of the larva itself.

### 4.5 Statistical Analysis

Experiments were conducted with three pairs of non-related male–female crosses. The Chi-square test was employed to compare the distribution of swimming larvae in response to 5-HT and DA with the control. The chi-square test calculations were performed using an interactive tool for chi-square tests of goodness of fit and independence (https://www.quantpsy.org/chisq/chisq.htm). The Spearman correlation test was conducted using XLStat v. 2016.02.28451. p-values less than 0.05 were considered statistically significant. Variances were calculated in double-centered samples using Microsoft Excel 2016. Graphs were generated with GraphPad Prism v. 8 and Microsoft Excel 2016, and mean ± SEM values are provided.

## 5 Author Contributions

Study concept and design — EGI, EEV, and ALO; collection of biological material — ALO, MYK, MNS; data acquisition — ALO, EGI, and EEV; data analysis and interpretation — EGI, ALO, and EEV; drafting of the manuscript — ALO, EEV, and EGI; resourses and funding – VVS; administrative and material support — EEV. All authors have read, edited, and agreed to the published version of the manuscript.

## 7 Informed Consent Statement

Not applicable

## 8 Data Availability Statement

Not applicable

## 9 Acknowledgments

The authors are grateful to Prof. Alexander Ereskovsky, Dr. Salim Daytov and members of laboratory of embryology NSCMB FEB RAS for the help with the larvae obtaining and culture. We are grateful the staff of the Vostok Biological Station (NSCMB FEB RAS), Far East, Russia, and Marine Station of Endoume, Mediterranean Institute of marine and terrestrial Biodiversity and Ecology (Institut Méditerranéen de Biodiversité et d’Ecologie marine et continentale (IMBE), Station Marine d’Endoume), Marseille, France. The research was done using equipment of the Core Centrum of the Institute of Developmental Biology RAS (IDB RAS RP # 0088-2021-0020), the Center for molecular and cell technologies, Center for Culturing Collection of Microorganisms, center “CHROMAS” of Saint Petersburg State University and “Taxon” Research Resource Center of Zoological Institute RAS,and resource center “Observatory of Environmental Safety”, Research Park of St. Petersburg State University. The experimental research with 5-HT and DA effects on the larvae was conducted in the frame of RSF, grant № 22-14-00375.

## 11 Institutional Review Board Statement

Not applicable

---

**Figure S1.**
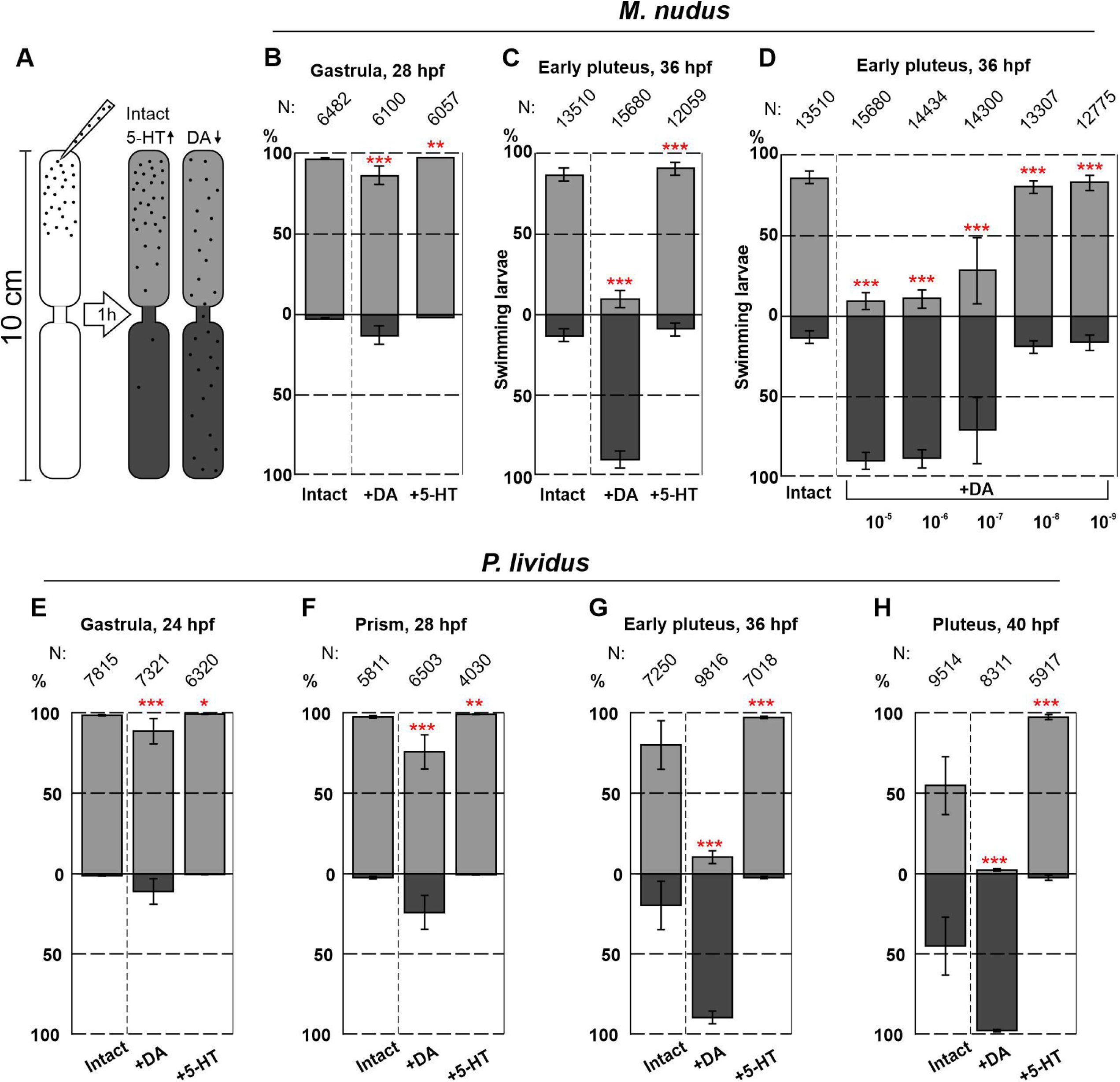
A Preliminary Behavioral Assay to Assess the Impact of DA and 5-HT on Larval Swimming Ability at Different Ages. (A) presents a schematic representation of the experimental setup. (B, C) depict the numbers of upward and downward swimming gastrula and early pluteus larvae of *M. nudus* in response to DA and 5-HT. (D) demonstrates that the downward swimming of *M. nudus* early pluteus in response to DA is dose-dependent, with 1 µM being the minimal concentration required to affect swimming. (E-H) show that the application of 5-HT results in upward swimming in *P. lividus* larvae of all ages, while DA affects downward swimming ability stage-dependently. It’s noteworthy that the impact of DA increases with larval age from gastrula to pluteus. N represents the number of larvae. Five independent replications for each test were performed. Chi-square test results are provided. Statistical significance: p < 0.05: *, p < 0.01: **, p < 0.001: ***.

**Figure S2.**
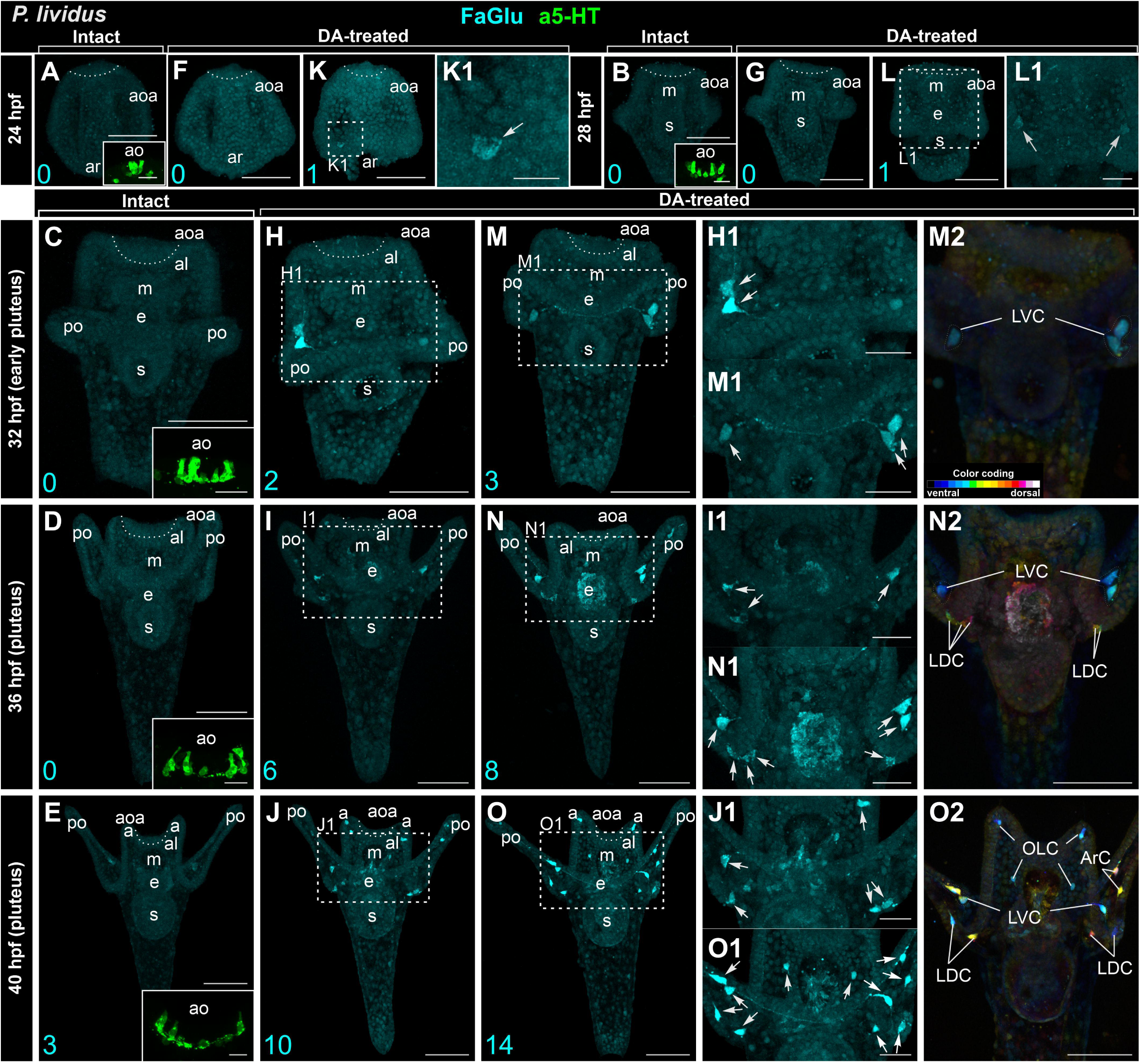
The Developmental Features of Dopaminergic and Serotonergic Neurons in *P. lividus* Larvae. DA-producing cells are visualized in intact larvae, while DA-uptaking cells are shown in DA-treated larvae (FaGlu, cyan), and 5-HT cells (a5-HT, green) are located in the apical organ. Panels (A, B, C, D, E) present a general view of DA-producing cells in *P. lividus* larvae at subsequent developmental stages from 24 hpf gastrula to 40 hpf pluteus. (A, B, C, D, E insets) provide a high magnification of 5-HT cells in the apical organ (ao). The figures highlight the gradual increase in DA and 5-HT cells with larval age. (F-K, G-L, H-M, I-N, J-O) offer a general view of DA-uptaking cells in larvae of subsequent developmental stages. Most of DA cells and their processes are associated with the arm. Note the variations in the number of DA cells, marked in cyan. (K1, L1) showcase a high magnification of the region with DA-uptaking cells in gastrula, where a maximum of two cells is visible. Representative samples of the minimal (H1, I1, J1) and maximal (M1, N1, O1) number of DA neurons in pluteus larvae are provided, with neurons marked by arrows. (M2, N2, O2) illustrate the relative position of DA cells in the larval body. Abbreviations; aoa – apical organ area; al – anterior lobe; ar — archenteron; ArC - arm cells; e — esophagus; LDC – lateral dorsal cells; LVC – lateral ventral cells; m — mouth; OLC – oral lobe cells; po — post-oral arms; s — stomach

